# A small fraction of progenitors differentiate into mature adipocytes due to constraints on the cell structure change

**DOI:** 10.1101/2021.03.10.434887

**Authors:** Mahmoud Ahmed, Trang Huyen Lai, Deok Ryong Kim

## Abstract

Differentiating 3T3-L1 pre-adipocytes are a mixture of non-identical culture cells. It is vital to identify the cell types that respond to the induction stimulus to understand the pre-adipocyte potential and the mature adipocyte behavior. To test this hypothesis, we deconvoluted the gene expression profiles of the cell culture of MDI-induced 3T3-L1 cells. Then we estimated the fractions of the sub-populations and their changes in time. We characterized the sub-populations based on their specific expression profiles. Initial cell cultures comprised three distinct phenotypes. A small fraction of the starting cells responded to the induction and developed into mature adipocytes. Unresponsive cells were probably under structural constraints or were committed to differentiating into alternative phenotypes. Using the same population gene markers, similar proportions were found in induced human primary adipocyte cell cultures. The three sub-populations had diverse responses to treatment with various drugs and compounds. Only the response of the maturating sub-population resembled that estimated from the profiles of the mixture. We then showed that even at a low division rate, a small fraction of cells could increase its share in a dynamic two-populations model. Finally, we used a cell cycle expression index to validate that model. To sum, pre-adipocytes are a mixture of different cells of which a limited fraction become mature adipocytes.

## 1 Introduction

Cell cultures contain populations of non-identical cells. Early on, pre-adipocytes cell cultures were recognized to be heterogeneous. Green *et al.* described the heterogeneous nature of differentiating 3T3-L1 cell [1]. Lee *et al.* showed that adipocyte progenitors within a single fat depot give rise to functionally distinct adipocytes [2]. Cell cultures never attain a 100% differentiation efficiency; thus, only a fraction of the preadipocytes are responsive. A key question is how large or small this fraction is. Loo *et al.* showed that a minority of pre-adipocytes gives rise to mature adipocytes with characteristic gene expression and lipid droplets [3].

The varying cellular response to the induction media could be because of their intrinsic properties or is a stochastic process. Stochasticity and cell-cell interactions were suggested as key contributors to the differentiation of stem cells [4]. If the pre-adipocytes are intrinsically different, we should be able, in principle, to find mechanisms that confer responsiveness/resistance to the sub-populations. A study proposed that the adipocyte’s ability to mature and accumulate fat is heritable. Modifying the key adipogenic transcription factor peroxisome proliferator-activated receptor gamma (*Pparg*) promoter region marks the cell capacity to differentiate, and re-differentiate [5].

Differentiation is a transcriptional and morphological development event. Cells proliferate only for one day after the 3T3-L1 induction in what is known as post-confluent mitosis followed by growth arrest [6]. Cell division and DNA replication are necessary to allow transcription factors into the open chromatin. This, in turn, drives the development of the new phenotype. Within days, the cell accumulates lipids in droplets and remodels its interior into a distinct fat cell.

Changes in the cytoskeleton and extracellular matrix rarely feature during the study of adipocyte differentiation. This is unfortunate given their prime importance to the morphological development of the adipocytes. It is clear from research done in stem cells with differentiation potentials; they have to respond to extracellular signals, not only by modulating the gene expression but their internal structure [7, 8]. Yang *et al.* showed that remodeling the cytoskeleton through the acetylation of alpha-tubulin is necessary for the differentiation of pre-adipocytes [9]. Moreover, it was even suggested that actin cytoskeleton might interfere with the transcriptional regulation of the differentiation process by controlling the translocation of antagonistic nuclear factors [10].

Profiling by RNA-Seq is an efficient way to study gene expression changes between two or more conditions. However, the expression signal, in this case, is averaged over a mixed population of various cell types. Single-cell RNA-Seq (scRNA-Seq) is the most straightforward way to test the gene expression at the level of the individual cells. Because these datasets are not always available, deconvolution of bulk RNA-Seq data can be an alternative. In this study, the subpopulations in the mixture were estimated from their gene signatures and studied separately.

Convex analysis of mixtures (CAM) is an unsupervised method to deconvolute the profiles of samples from mixed phenotypes into their constituents [11]. The creators of the method applied it to a dataset of gene expression in yeast at different cell cycle stages. CAM could detect a composite of sub-populations consistent with the true labels. In addition, the sub-populations gene markers identified by CAM were related to the corresponding phases. Herrington *et al.* used this method to derive distinct phenotypes in arterial tissues. Classification of the cell types based on their markers was consistent with tumor necrotic factor (TNF) alpha pathway activation [12].

Here we describe deconvoluting the gene expression of differentiating 3T3-L1 pre-adipocytes from bulk RNA-Seq data. The goal is to determine the distinct phenotypes in the cell cultures and estimate their fractions. Then we showed which of these fractions responded to the induction stimuli and became mature adipocytes. Finally, we attempted to explain the differences between the sub-populations as to which functions their specific markers involve in, which type of cells they give rise to at the end of the differentiation course, and their response to treatment with various drugs and compounds.

## 2 Materials & Methods

### 2.1 Cell culture & differentiation

3T3-L1 is a mouse pre-adipocyte cell line that can be induced to differentiate into mature adipocytes when treated with a chemical cocktail. A chemical cocktail of 1-Methyl-3-isobutylxanthine, Dexamethasone and Insulin (MDI) is known to induce the differentiation [1]. The treatment starts with a fully confluent 3T3-L1 cell culture (preadipocyte), accumulating lipid into lipid droplets within days. The time course comprises three stages; day 2–4, early; up to day 7 intermediate; and up to 14 days is the late-differentiation stage. A similar protocol produces a comparable differentiation course in human primary preadipocytes.

### 2.2 Data curation & processing

#### 2.2.1 RNA-Seq data

We surveyed GEO and SRA repositories for high-throughput sequencing data of MDI-induced 3T3-L1 and human primary pre-adipocyte samples at different time points (Table 1). Ninety-eight RNA-Seq samples were included in the study. This dataset was previously curated, and the processed data was packaged in a Bioconductor experimental data package `CuratedAdipoRNA` [13]. Briefly, raw reads were downloaded from the SRA ftp server using `Wget`. FASTQC was used to assess the quality of the raw reads [14]. For RNA-Seq, raw reads were aligned to UCSC mm10 mouse genome using HISAT2 [15]. `FeatureCountS` was used to count the aligned reads (bam) in known genes [16].

**Table 1:** RNA-Seq datasets of differentiating 3T3-L1 cells.

| SRA ID | Time (hrs) | Ref. |
| --- | --- | --- |
| SRP010905 | 192:192 | [22] |
| SRP029602 | -48:144 | [23] |
| SRP029985 | 0:168 | [24] |
| SRP033778 | -48:240 | [25] |
| SRP041765 | 0:4 | [26] |
| SRP045772 | 0:48 | [27] |
| SRP051775 | 168:168 | [28] |
| SRP066991 | -96:168 | [29] |
| SRP078506 | 0:48 | [30] |
| SRP090166 | 0:168 | [31] |
| SRP092793 | 240:240 | [32] |
| SRP100171 | 0:192 | [33] |
| SRP100871 | 0:168 | [34] |
| SRP102079 | 0:4 | [35] |
| SRP109255 | -48:24 | [36] |
| SRP119259 | 192:192 | [37] |

#### 2.2.2 Microarray data

We conducted a similar survey of the literature for studies that subject the same cell line model for chemical or genetic perturbations (Table 2). Forty-two microarray of mature adipocytes (> 7 days post MDI-induction) treated with thiazolidinediones (TZD), four short-chain fatty acids (SFA), or three nonsteroidal anti-inflammatory drugs (NSAID) were previously collected and curated. The processed data were packaged in another experimental data package `curatedAdipoArray` [17]. In addition, 24 microarray samples of MDI-induced human primary pre-adipocytes were downloaded from GEO in the form of processed probe intensities (GSE98680) [18].

**Table 2:**
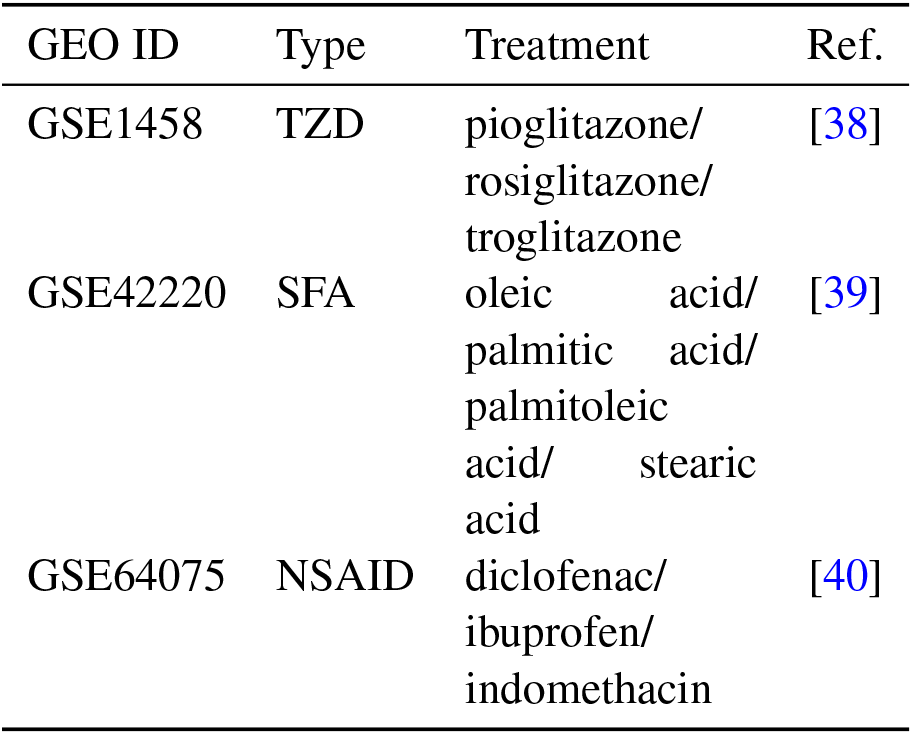
Microarray datasets of differentiating 3T3-L1 under pharmacological perturbations.

| GEO ID | Type | Treatment | Ref. |
| --- | --- | --- | --- |
| GSE1458 | TZD | pioglitazone/<br>rosiglitazone/<br>troglitazone | [38] |
| GSE42220 | SFA | oleic acid/<br>palmitic acid/<br>palmitoleic<br>acid/ stearic<br>acid | [39] |
| GSE64075 | NSAID | diclofenac/<br>ibuprofen/<br>indomethacin | [40] |

Microarray data were downloaded from GEO in the form of probe intensities. Probe metadata (gene symbols) were used to collapse the intensities into gene level using the maximum average [19]. Data were normalized using normalize between arrays, and batch effects were removed using empirical Bayesian adjustments [20, 21].

### 2.3 Identifying adipocyte populations

Convex Analysis of Mixtures (CAM) was used to deconvolute gene counts expression profiles [11]. This analysis was applied using `debCAM` [41]. Briefly, the scatter simplex of the cell mixture is a rotated and compressed version of the scatter simplex of the pure sub-populations. The sub-populations and their markers reside at the extremities of the shape. The one-vs-everyone fold-change was calculated for all genes and used to select gene markers. Using the expression of the markers, the relative fractions of the subpopulations in the mixture were estimated by standardized averaging. The estimated fractions were used to deconvolute the mixed expressions into sub-populations specific profiles by least-square regressions.

### 2.4 Validating markers & populations

A supervised version of CAM was applied using different sets of markers and cell profiles. To isolate the effect of the gene markers’ set size, the fractions of the sub-populations were re-estimated using a smaller set of the identified markers. The set size was chosen to be equal to that of the smallest sub-populations. To isolate the effect of outliers, the same process was repeated multiple times using a set chosen at random from the identified markers. The analysis was applied to 3T3-L1 transcription activity (POLR2A occupancy) and the expression profiles of human primary adipocytes using the full set of markers.

### 2.5 Gene set over-representation

Lists of sub-populations specific markers were annotated using the gene ontology terms (`org.Mm.eg.db`) and tested for overrepresentation using `ClusterProfiler` [42, 43]. For each gene set, the observed fraction of gene in the list was compared to the fraction expected by chance. P-values were calculated for each gene set and adjusted for multiple testing using false discovery rate (FDR).

### 2.6 Differential expression analysis

The effect of drugs and compounds treatment on gene expression of mature adipocytes was evaluated at the mixture and population levels. The differential gene expression between treatment and control groups was calculated using `LIMMA` [20]. Population-specific gene expression analysis was applied to the same comparisons using `PSEA` [44]. The previously identified markers of the three sub-populations were used to construct a reference population and perform one-to-one comparisons. P-values were calculated for each gene and adjusted for multiple testing using FDR.

### 2.7 Software & reproducibility

The analysis was conducted in an R and using Bioconductor packages [45, 46]. The software environment is available as a docker image (https://hub.docker.com/r/bcmslab/adipo_deconv). The code for reproducing the figures and tables in this document is open-source (https://github.com/BCMSLab/adipo_deconv).

## 3 Results

### 3.1 Pre-adipocytes cell cultures consist of three distinct sub-populations

Averaging the gene expression in a given sample reflects the dominant phenotypes of cells during the course of differentiation. Pre-adipocytes were dissimilar to early and late-differentiated cells. We used as a measure of dissimilarity the distances among the samples within and in-between stage. The distances within the pre-adipocytes samples differed from early (*t* = 24.07; *P* < 0.01) and late-differentiated (*t* = 30.05; *P* < 0.01) samples. Moreover, the stage of differentiation explained about half the variance among the samples (Figure 2A). Un-differentiated and differentiating cells separated across the first two dimensions (*k* = 2; goodness-of-fit > 0.47) of the pairwise distances of the corresponding samples.

**Figure 1:**
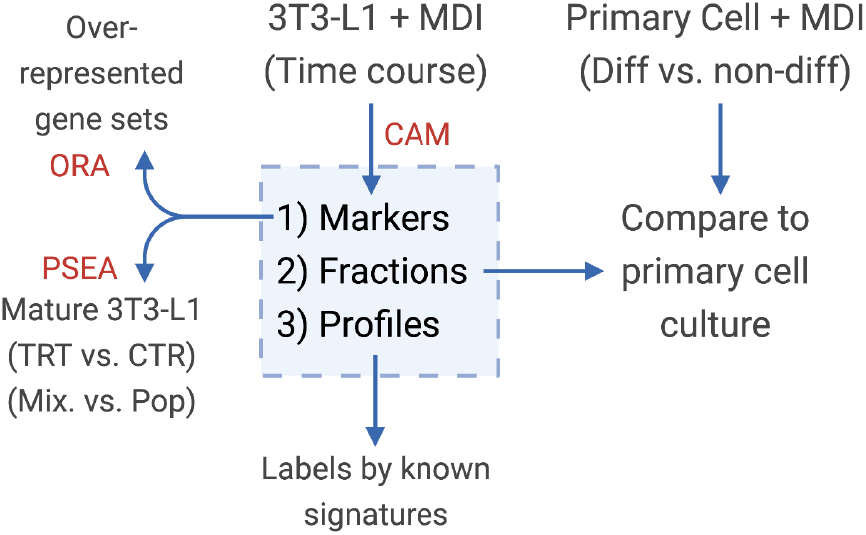
Diagram of the study design. Convex analysis of mixtures (CAM) was applied on MDI-induced 3T3-L1 time-course gene expression to estimate the sub-population 1) markers 2) fractions, and 3) specific expression profiles. Markers were tested for over-representation in gene sets (ORA). Sub-population fractions were validated in primary adipocytes, and expression profiles were examined for known gene signatures.

**Figure 2:**
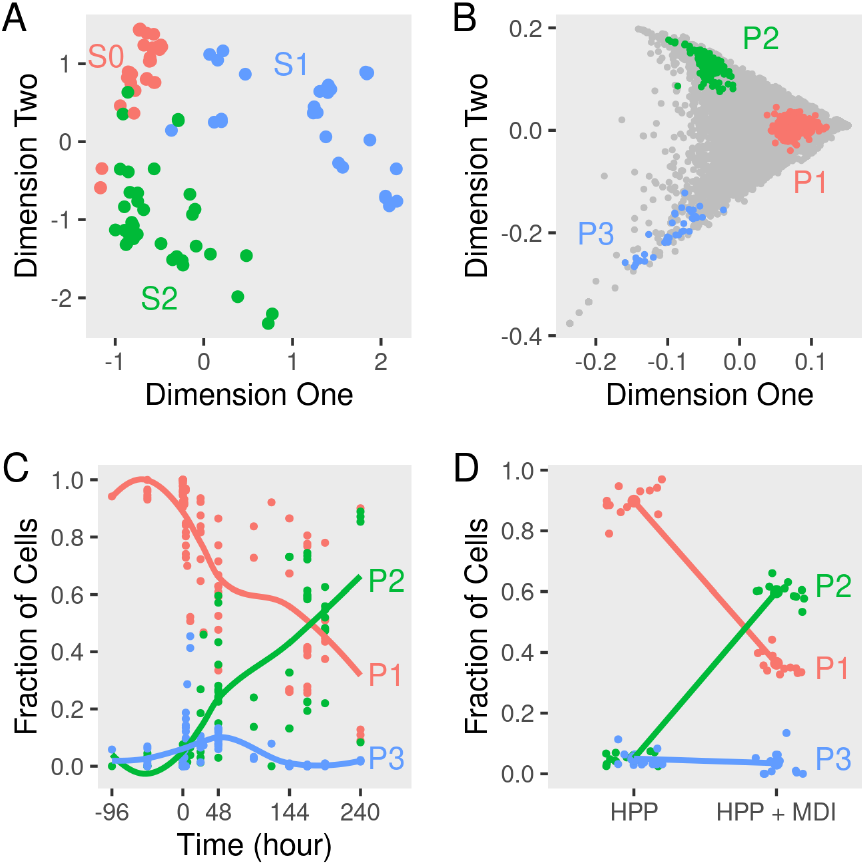
Dimensions reduction and deconvolution of differentiating adipocytes gene expression. Gene expression of MDI-induced 3T3-L1 pre-adipocytes (n = 98) and human primary adipocytes (HPP) (n = 24) was determined by RNA-Seq or microarrays at different times. (A) Euclidean distances among samples were calculated from the transformed gene counts. Samples were laid out in two dimensions to approximate the distances and colored by stage (S0, S1, or S2) for non-induced, early or, late differentiation. (B) The scatter simplex of the cell mixture is a rotated and compressed version of the scatter simplex of the pure sub-populations. In two dimensions, the sub-populations (P1, P2 & P3) and their markers reside at the extremities. (C) Using these markers, the relative fractions of the sub-populations in the mixture of differentiating 3T3-L1 were estimated by standardized averaging. (D) The relative fractions of the sub-populations in the mixture of differentiating HPP were estimated by standardized averaging of the same markers.

The 3T3-L1 cell cultures before and after the induction of differentiation consist of mixtures of cells. Average expression suffices to study the changes in gene expression over time, but the signal of each gene is potentially a composite of its expression in two or more cell types. These may differ in their phenotypes and/or genotypes. At each stage/time point of differentiation, the composition of the culture may differ, and the fractions of the sub-populations may change. Deconvoluting the gene expression of differentiating adipocytes would help address these points.

To determine the sub-populations in the cultures, we applied the convex analysis of mixtures (CAM) to the mixed gene expression profiles of differentiating adipocytes. The scatter simplex of the expression of the mixture is a transformed version of that of the pure subpopulations. When projected in low-dimensions, we observed three distinct archetypes characteristic of at least three groups of cells (Figure 2B). Gene markers of these sub-populations were identified as genes that are differentially expressed in one sub-populations vs. others (fold-change > 1; bootstrapped lower bound confidence interval > 0) (Figure 3A). Markers were used to estimate the relative fractions and expression profiles of the subpopulations in each sample.

**Figure 3:**
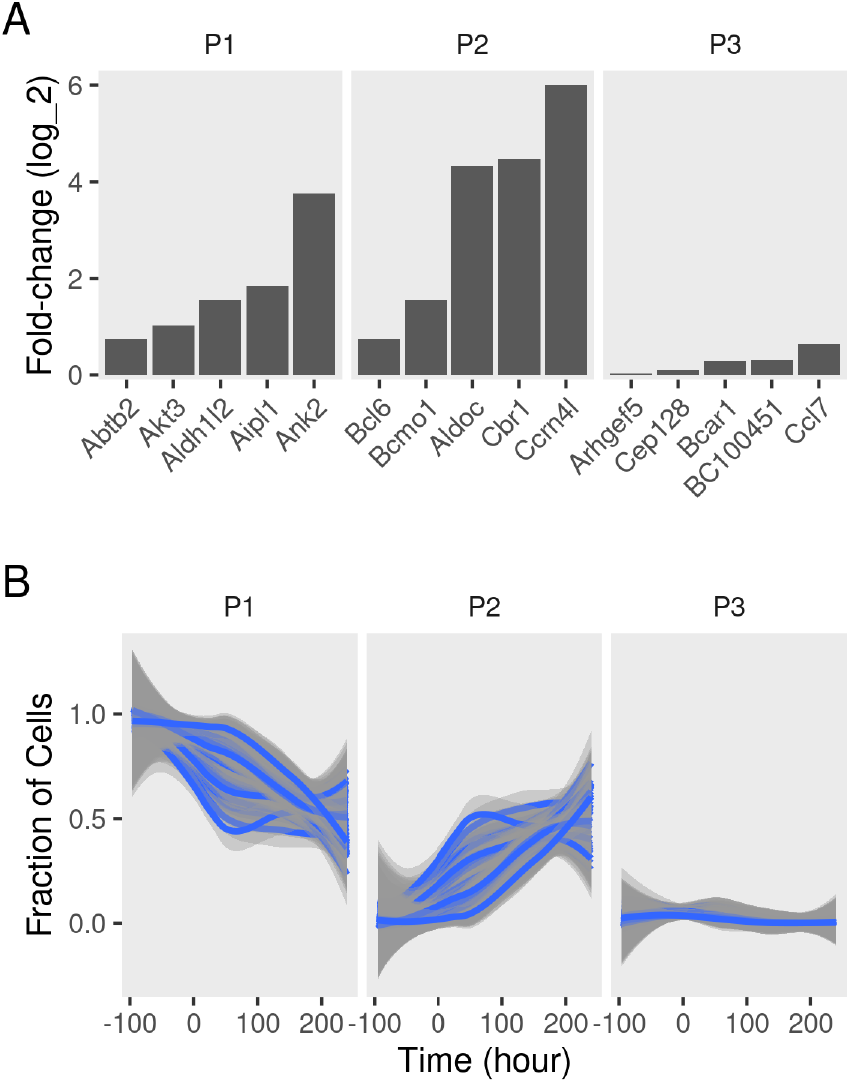
Sub-populations gene markers and fractions in differentiating adipocytes. Gene expression of MDI-induced 3T3-L1 preadipocytes (n = 98) was determined by RNA-Seq at different time points. Convex analysis of mixture was used to estimate the relative fractions of sub-populations cultures and their specific expression profiles. (A) The one-vs-everyone foldchange was calculated for all genes. The foldchange of the top 5 genes in each sub-populations (P1, P2 & P3) is shown. (B) Sets for subpopulation markers (set size = 35; iteration number = 100) were re-sampled from the originally identified gene markers. The markers sets were used to re-estimate the relative fractions of the sub-populations (P1, P2 & P3) in the mixture were by standardized averaging. Average fractions are shown as smoothed lines (blue), and the standard errors are shown as ribbons (gray).

Various sets of sub-population markers gave patterns not very different from the above. We chose random sets of markers of the same size for the three sub-populations (bootstrapping). Lowering the number of markers and randomly resampling from the original set gave similar estimates of the fractions of the three populations at most time points (coefficient of variation < 10%; bias < 0.1) (Figure 3B).

### 3.2 Mature adipocytes originate from a small fraction of pre-adipocytes

The starting cell culture is composed of disproportionate fractions of phenotypes that change with time (Figure 2C). One of the three groups (P1) makes up over 95%, and the rest is a mixture of the two other groups. Over the course of differentiation, the fraction of cells of the dominant group decreases to less than 40% of the late samples. In response to the induction stimulus, two factions of cells increase their share of the culture initially. Only one group (P2) continues to increase in size until the end and reaches more than 60% of the cells.

The three sub-populations were identified based on their specific gene expression profiles. Adipogenic and lipogenic genes marked the second sub-population (P2) as the ones responding to the induction stimulus and forming mature cells (Figure 4A). The first group of genes included the transcription factors *Pparg* and *Cebpb* involved in regulating the differentiation process. Those were highly expressed in P2 compared to the two other sub-populations (fold-change > 7; lower confidence interval > 0). The second group of genes code for key enzymes such as *Fasn* and *Lpl* in the lipid synthesis pathway and are essential for the accumulation of lipid. Those were also expressed at a higher level in P2 (fold-change > 1.5; lower confidence interval > 0).

**Figure 4:**
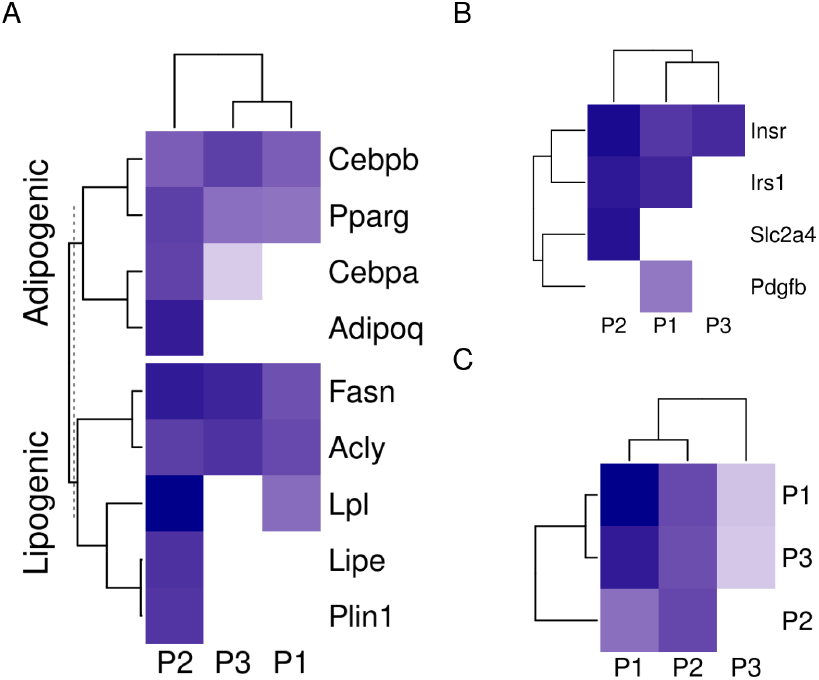
Sub-population specific gene expression in differentiating adipocytes. Gene expression of MDI-induced 3T3-L1 pre-adipocytes (n = 98) was determined by RNA-Seq at different time points. Convex analysis of mixture was used to estimate the relative fractions of subpopulations in cultures. The estimated fractions were used to deconvolute the mixed expressions into sub-populations specific profiles by leastsquare regressions. (A) The expression of adipogenic and lipogenic gene markers is shown in three sub-populations. (B) The expression of insulin signaling genes is shown in the three sub-populations. (C) Spearman’s rank correlation coefficient (*r_s_*) among the gene expression in the sub-populations of differentiating 3T3-L1 and primary human adipocytes (HPP) were calculated and shown as color values (white, low; blue, high). Sub-populations are indicated as (P1, P2 & P3) and expression as color values (white, low; blue, high).

The differences in gene expression between the two sub-populations extended to more than adipogenic and lipogenic related genes. Insulin metabolism-related genes were different expressed in the differentiating sub-population (P2) compared to the other two (fold-change > 3; lower confidence interval > 1.7) (Figure 4B). Although insulin receptor (*Insr*) and its substrate (Irs1) were expressed in the two major subpopulations, the glucose transporter solute carrier family 2 member 4 (*Slc2a4*) was expressed in P1 only. Platelet-derived growth factor subunit B (*Pdgfb*) exclusive expression in P2 indicates non-responsive cells to glucose.

A similar mixture of sub-populations constitutes the cell culture of human primary preadipocytes. The fractions of cells in primary preadipocytes corresponding to the previously identified markers resembled that that were estimated from 3T3-L1 cells (Figure 2D). Similarly, one of the sub-populations (P2) grows to dominate the final culture of MDI differentiated cells. The rest (P1 and P3) did not or only briefly responded. The specific gene expression profiles of the subpopulation in the primary cells were similar to that in the mouse cell model (Figure 4C). Because expression profiles come from two different types of cell lines, the expression of the sub-populations in each were strongly correlated (Spearman’s rank correlation coefficient (*r_s_*) > 0.6; *P* < 0.001). At the same time, the corresponding sub-populations were strongly correlated between the two cell lines. This was the case for P1 (*r_s_* = 0.47; *P* < 0.001) and P2 (*r_s_* = 0.4; *P* < 0.001). The only exception was the weaker correlations of P3 (*r_s_* = 0.2) which was less than that of the mouse P3 with human P1 (*r_s_* = 0.4; *P* < 0.001).

### 3.3 Sub-populations of mature adipocytes respond discrepantly to various compounds

If, in fact, the mature adipocytes are a composite of different sub-populations, we expect to see a heterogeneous response to the treatment of different compounds. We collected data from MDI-induced adipocytes after day 7 and treated them with various compounds. These compounds included three thiazolidinediones (TZD), four short-chain fatty acids (SFA), and three nonsteroidal anti-inflammatory drugs (NSAID). The response of the cell mixture was evaluated by the regular differential expression, and that of the sub-populations was evaluated by population-specific expression analysis (PSEA). By comparing the response in the adipocyte mixture to that of the individual sub-populations, we could identify the effect of each compound on the mature adipocytes specifically.

We found that compounds from the same family elicited similar responses as expected (Figure 5A). The response to TZD: rosiglitazone, pioglitazone, troglitazone on the average gene expression was highly correlated (Pearson correlation coefficient (*r*) > 0.5; *P* < 0.001). Similarly, the fold-change in response to SFA and NSAID treatment was correlated (*r* > 0.4 and *r* > 0.6; *P* < 0.001). Only small correlations were found between the fold-change response between groups of compounds. TZD treatments were positively correlated with the NSAID and negatively correlated with that of SFA. NSAID and SFA were negatively correlated.

**Figure 5:**
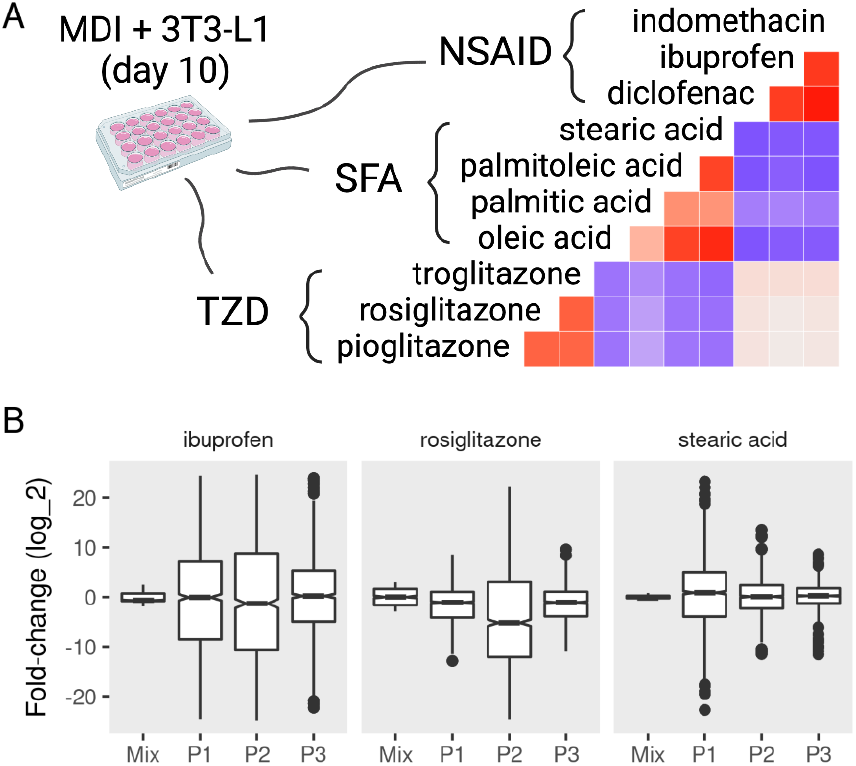
Mixture and sub-population specific response of mature adipocytes to pharmacological perturbations. MDI-induced 3T3-L1 mature adipocytes (> 7 days) were treated with dimethyl sulfoxide (DSMO), nonsteroidal antiinflammatory drugs (NSAID), short-chain fatty acids (SFA), or thiazolidinediones (TZD). Gene expression of the samples (n = 42) was determined by microarrays. Previously identified subpopulations (P1, P2 & P3) specific gene markers in the 3T3-L1 cells were used to estimate the subpopulation specific expression profiles. A) The correlations between the fold-change in response to treatments are shown as color values (blue, negative; red, positive). The distribution of the foldchange of the top 500 differentially expressed genes in the mixture and sub-populations in response to rosiglitazone, ibuprofen, and stearic acid.

The response on the sub-population level to the treatment of different compounds often varied from the mixture and among each other (Figure 5B). Treatment with rosiglitazone produced more pronounced differential expression in P2 compared to the other populations (KS test *D*^−^ < 0.15; *P* < 0.001) as well as the mixture (*D*^−^ < 0.24; *P* < 0.001). A similar pattern was observed with the treatment of ibuprofen (*D*^−^ < 0.13; *P* < 0.001). By contrast, P1 was more different when adipocytes were treated with stearic acid (*D*^−^ = 0.43; *P* < 0.001).

### 3.4 A dynamic model of two subpopulations in a fully confluent cell culture

To show the validity of these observations, we constructed a model of sub-populations dynamics and compared it with further observations from the data. In this model, two sub-populations of different sizes grow at different rates in response to the environment in a fully confluent cell culture (Figure 6A). *P*2 is a small sub-population that grows at a rate *r, rP*1 while *P*1 is a large sub-population that decreases in size at a smaller or the same rate –*r*, –*rP*1. In this case, the total population size (*N*) can be estimated as (*P*2 – *P*1)*r*. This predicts that the total population (*N*) should decrease initially and bounces to the same size or higher after two days, given a small *r*. This system can be summarized as follows:

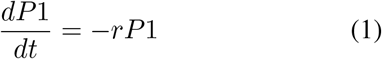

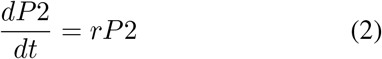

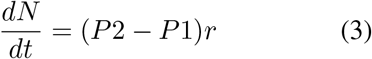

**Figure 6:**
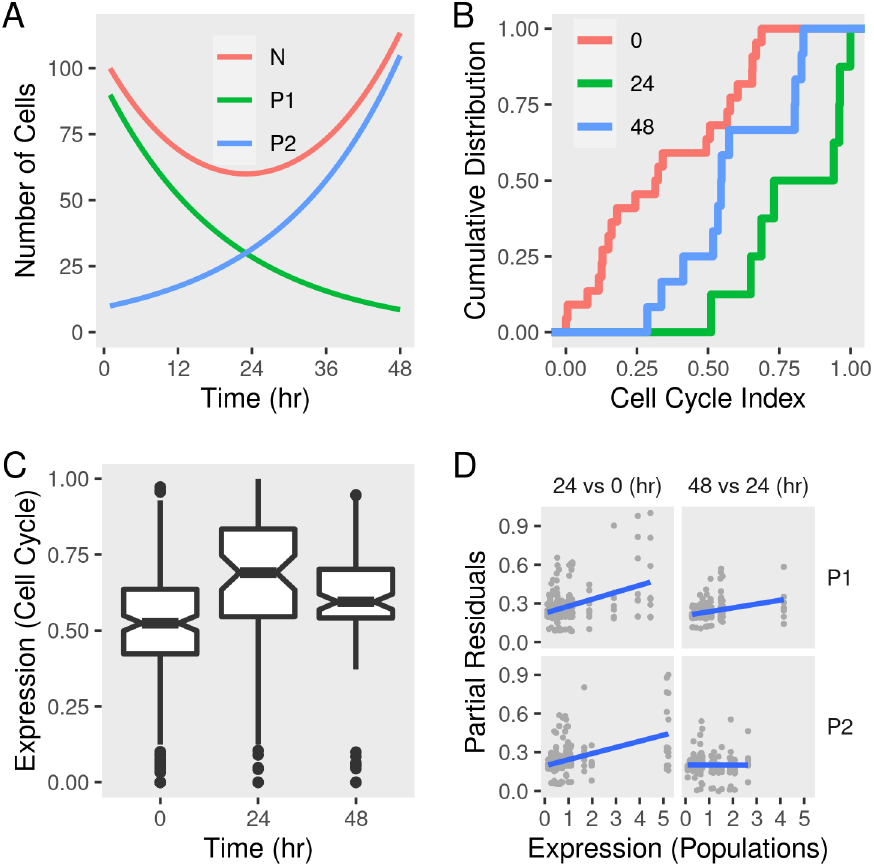
Sub-population dynamics and cell cycle gene expression signature in differentiating adipocytes. (A) A toy model of two subpopulations P1 and P2 growing at rates – *r* and *r*, respectively and the total population *N*. The change in the total number of cells and each subpopulation was calculated over time given reasonable initial states and parameter values. (B) Gene expression of MDI-induced 3T3-L1 preadipocytes (n = 42) was determined by RNA-Seq at three time points. The cumulative distribution function of eight-cell cycle genes in differentiating adipocytes at 0, 24, and 48 hours. (C) The distribution of the expression of eightcell cycle genes in differentiating adipocytes at the same time points. (D) Previously identified sub-populations (P1 & P2) specific gene markers were used to estimate the sub-population specific expression profiles. Regression models were used to estimate the differential expression between 24 and 0 hours and between 48 and 24 hours. Partial residuals of each comparison and populationspecific expression are shown.

A fully confluent cell culture imposes a constraint on the rate at which the sub-populations can grow or die (*r*). We chose a value that reestablishes the population size within a reasonable period (two days). The rate at which one population can grow is bound by the rate at which the size of the other population shrinks. The values of the initial states (*P*1, *P*2, and *N*) were suggested by the previous analysis where the growing subpopulation represents a small fraction of the total at the start of the cell culture.

### 3.5 Cell cycle index fits the dynamics of two sub-populations

We used a cell cycle gene expression signature proposed by Dolatabadi *et al.* to estimate the cell cycle index at population and sub-population levels [47]. We found that, adipocytes at day 1 had higher cell cycle indecies than pre-adipocytes (KS test *D*− = 0.7; *P* < 0.01) and adipocytes at day 2 (*D*^−^ = 0.5; *P* > 0.05) (Figure 6B). This pattern reflects the higher expression of cell cycle genes at day 1 compared to pre-adipocytes (enrichment score = 0.7; *P* < 0.05) and adipocytes at day 2 (enrichment score = 0.6; *P* > 0.05) which indicates a larger proportion of the cells in G0/S phase and, therefore, less active divisions (Figure 6C). This was consistent with the predicted population size of the toy model.

The sub-population specific differential expression confirmed that the two populations contribute differently to the expression of cell cycle genes and therefore to the active cell division (Figure 6D). This is clear from the correlation between the expression of the cell cycle genes and the reference (population signals). It is not clear from this analysis how to assign specific values to each sub-population at each of the two comparisons. This is because the expression is calculated for each of the eight genes individually. Moreover, the index reflects a continuous transition from G1 to S phase of the cell cycle rather than discrete categories. This would be affected by the size of the population at different time points and, therefore, their contribution to the overall expression of the genes constituting the index.

### 3.6 Resistance to differentiation may be due to structural restrictions on the cell and/or commitment to other lineages

Several functional sets of genes were overrepresented (count ≥ 3; false-discovery rate < 0.2) in the group of markers of the growing subpopulation (P2). These were related to fat differentiation, nuclear receptors and, transcriptional activities (Table 3). More specifically, many of the characteristic markers were part of the process of *brown fat cell differentiation.* Molecular functions such as *nuclear receptor* and *steroid hormone activities* were over-represented in the same group. Several of these markers localized to cellular components such as *transcription factor complex*. Alternatively, the phenotype that did not show these characteristics could be thought of as resisting the induction stimulus. Indeed, completely different processes and functions were over-represented by the markers of the other two populations.

**Table 3:**
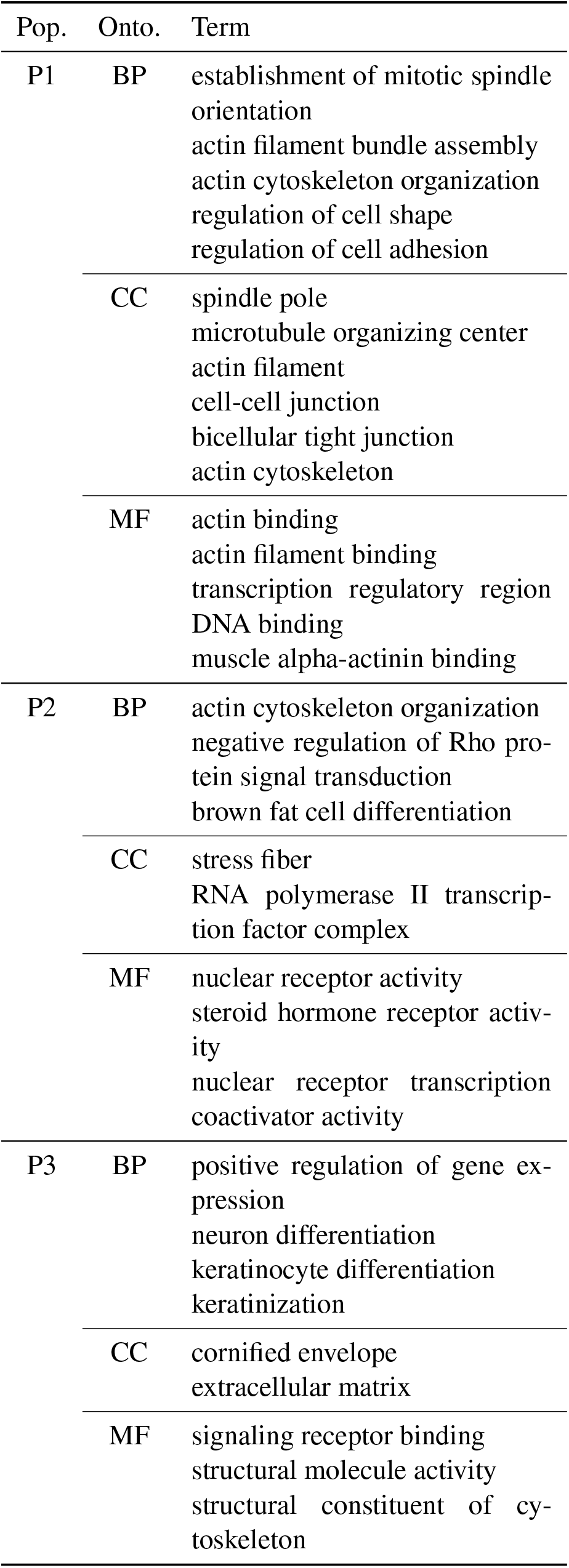
Gene set over-representations in the sub-populations specific gene markers.

The markers of the other sub-populations were enriched in terms related to cellular structural processes and components (Table 3). The first group (P1) had multiple markers related to *mitotic spindles*, *microtubules*, *actin* and *cytoskeleton*. All are essential components of the cell structure and necessary for allowing morphological changes. In addition to these terms, several processes related to the assembly and organization of these components were enriched by P1 markers.

The third group (P3) had terms related to structural components, namely the *extracellular matrix* and molecular functions *structural constituent of cytoskeleton* enriched by its markers. But also biological processes terms related to the differentiation of cell types other than adipocytes, *neurons* and *keratinocytes differentiation* were overrepresented in the P3 set of markers.

### 4 Discussion

We found that the pre-adipocyte cell culture is composed of three distinct sub-populations. After the differentiation induction, a minority of cells grows and transforms into mature adipocytes. This fraction of cells have a gene expression profile conducive to activities related to the nuclear receptors, fat differentiation, and gene transcription. The sub-populations of cells that do not respond to induction are under structural constraints or are committed to differentiate into other cell types. Figure 7 summarizes these observations in a graphical representation, which we further discuss below.

**Figure 7:**
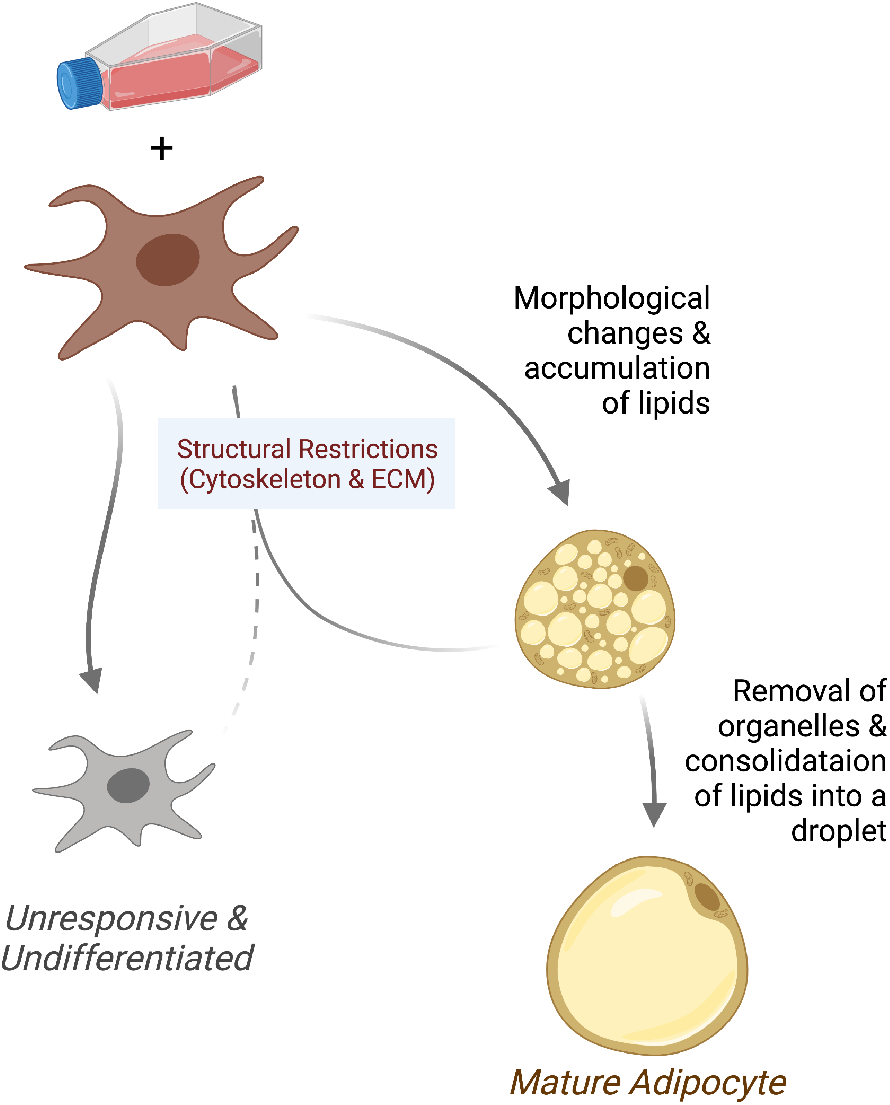
Model of the differentiating adipocyte sub-populations. A small fraction of pre-adipocyte fibroblasts growing in differentiation media responds to adipogenic stimuli. Responsive cells undergo morphological changes and accumulate lipids into small droplets. Adipocytes reach maturity after intracellular organelles are removed and fat consolidated into bigger droplets. Cells that fail to overcome structural restrictions such as those in the cytoskeleton and extracellular matrix do not undergo morphological changes and remain undifferentiated.

Identifying the factors that render fibroblasts responsive or resistant to the induction medium would help achieve better differentiation efficiency. More importantly, it would improve our understanding of adipose tissue composition and how to encourage or discourage the terminal maturation of the adipocytes.

In the context of CAM, exclusively enriched genes in a subset of cells are the markers for the set [11]. The expression of other genes (coexpressed) is the combination of their expression in the sub-populations. The expression of all genes in the mixture forms a k-dimensional simplex with a telling geometry. The simplex shape encloses the data, and the sub-populations with their markers are at the extremes.

To demonstrate the validity of these findings, one has to show that 1) distinct populations can exist in the pre-adipocyte cell culture and that 2) it is possible for some cells but not others to increase their population size. Differences intrinsic to the cells, in the ability to respond to MDI, in the kind of cell they become, or a combination of the above could give rise to the observed pattern. Although the size of the sub-populations is dissimilar to start with, it only changes after the differentiation induction. Because only the fractions of each sub-population rather than their absolute size were estimated, the inflection is possible to achieve with either the duplication of the responsive cells or the shrinkage of the others. Indeed, pre-adipocytes can replicate after the induction [6]. Even mature adipocytes can divide and redistribute the lipid droplets between the daughter cells [48].

Previous studies pointed out the heterogeneous nature of adipocyte cell cultures and adipose tissue progenitors. Often these studies focused on the phenotypic and molecular differences among the identified populations. Most attribute these differences to causes in two categories: asynchronous differentiation and lineage commitment. The first refers to cells that are essentially similar but only take longer to differentiate [3]. Therefore, at any specific point, some cells would lack the characteristics of mature adipocytes. For example, poor-lipid accumulating cells would form larger droplets of lipid by merely prolonged simulation [49]. While our study does not rule out this possibility, it makes counter observations. First, the differences between the sub-populations are stark from the very beginning (pre-adipocytes). Second, the lagging sub-populations do not show any sign of differentiation either in adipogenic regulation or lipid synthesis. Finally, the dataset extends up to two weeks, twice the period of typical maturation.

Studies based on the adipose tissue progenitor cells suggest that some cells are either incapable of differentiating into mature adipocytes, originate in certain depots, or can only differentiate into other cell types (brown/beige) adipocytes. One study suggested that mouse visceral, but not subcutaneous, white adipose tissue arises from cells expressing Wilms tumor (*Wt1*) gene [50]. Another found that the adipocyte’s ability to accumulate fat is heritable [5]. Histone modification of PPARG promoter marked cells which can de-differentiate and re-differentiate. It’s even been shown that a minority sub-population lacking the typical adipocyte features might take on a regulatory role to influence its surroundings in a paracrine manner [51].

Here, we suggest that the mature adipocytes arise from a minority of the pre-adipocytes. Most pre-adipocytes cannot differentiate properly or accumulate lipids. This lack of responsiveness to the differentiation medium is either due to or associated with structural restraints. The major factors that differentiate the two types of cells are genes involved in the cellular cytoskeleton and actin filaments. Molecular differences also exist, most importantly in regulating adipogenesis, lipid synthesis, and insulin signaling.

## Acknowledgments

This study was supported by the National Research Foundation of Korea (NRF) grant funded by the Ministry of Science and ICT (MSIT) of the Korea government [2015R1A5A2008833 and 2020R1A2C2011416].

## Conflict of interest

The authors declare no conflict of interests.

## Notes

https://github.com/BCMSLab/adipo_deconv

### Competing Interest Statement

The authors have declared no competing interest.

